# Evolutionary turnover of protein structural disorder drives aberrant proteome remodelling in naked mole rat

**DOI:** 10.64898/2026.04.27.720964

**Authors:** Purva Mishra, Sukanya Bhattacharya, Jui Bhattacharya, Yachna Jain, Kuljeet Singh Sandhu

**Affiliations:** Department of Biological Sciences, Indian Institute of Science Education and Research (IISER) – Mohali, Knowledge City, Sector 81, SAS Nagar 140306

**Keywords:** Intrinsic protein disorder, Naked mole rat, Stress resistance, Aging, Redox-sensitivity, Tumour suppression, Proteostasis, Phase separation, Proteome remodeling

## Abstract

The naked mole rat is an evolutionary outlier among mammals, exhibiting extreme longevity, cancer resistance, hypoxia tolerance, pain insensitivity, eusociality, poikilothermy and other distinctive physiological traits, most of which likely resulted from its adaptation to highly adverse subterranean habitat. Despite accumulating data, the genetic and molecular basis underlying these traits remain poorly understood. Through analyses of 18 distinct protein attributes and allied datasets across hystricomorphs, myomorphs, carnivores, and primates, we observed lineage-specific evolutionary divergence in intrinsic protein disorder in the naked mole rat. The disorder turnover exhibited functional dichotomy. The gain of disorder preferentially associated with proteostasis, immune regulation, neurodevelopment, skeletal growth and tumour suppressive properties, while loss of disorder modulated mostly the cardiac development. The proteins that gained disorder in NMR exhibited lower degradation rates, consistent with stabilization through phase-separation, while the proteins losing disorder show pronounced divergence in gene expression. The disorder turnover was primarily driven by indels affecting functional regions including Pfam domains, ANCHOR-predicted binding sites, short linear motifs, stress induced modifications of Tyr, Met, and Cys residues. Notably, the gained disordered regions were inferred to be redox-sensitive, aligning to exceptional stress tolerance in naked mole rats. Collectively, our results highlight an unusual and previously overlooked large-scale proteome remodelling that drives the molecular evolution of extraordinary traits of naked mole rat.

**Graphical abstract:** 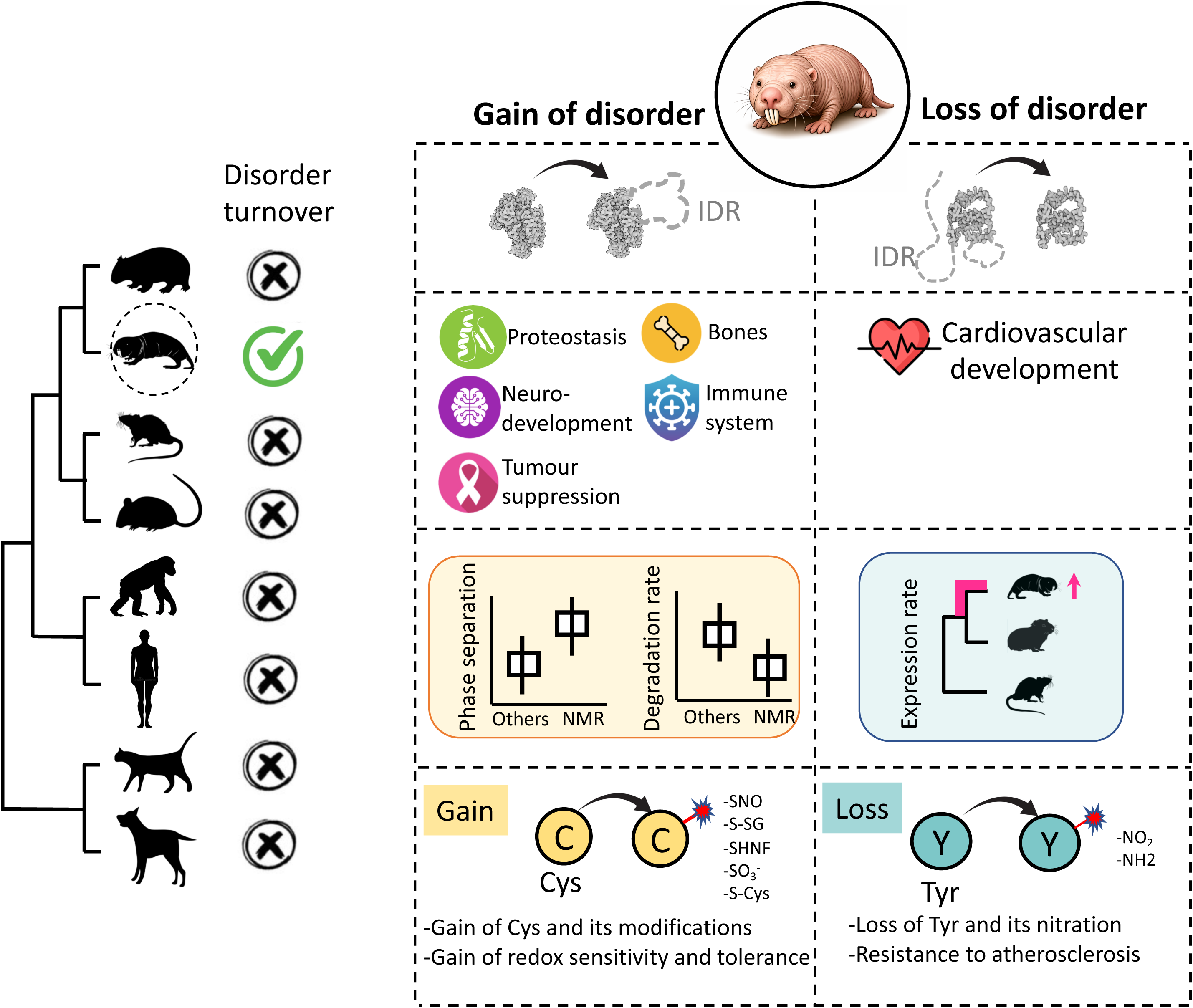

## Introduction

Naked mole rats (NMR) are extremophilic rodents that have evolved under hostile environmental pressures, most notably chronic hypoxia and hypercapnia in subterranean habitat^1–3^. To survive the strenuous conditions, NMR has developed a suit of extra-ordinary traits, including persistent poikilothermy^4^, slow metabolic rate^5–7^, enhanced resistance to oxidative stress^8–10^, elastic and wrinkled skin^11^, and dampened inflammatory response^12^. These adaptations may serve as foundation for species’ extreme longevity^13^. Though of similar size as mouse, which lives for 2-3 years, NMR can live up to 30 years^14,15^. Coherently, the species is resistant to age related disorders like cancer, cardiovascular abnormalities, immune senescence, neurodegeneration, and musculoskeletal loss^11,16^. Mechanisms such as early contact inhibition through high molecular weight hyaluronan, efficient DNA repair and dampened inflammatory response not only resist cancer, but also preserve cellular integrity over time^1117^. This is further complemented by their stable proteome and epigenome^1819^. Alongside Damara Mole Rat (DMR), they are the only known eusocial mammals, living in highly organized colony of around 70 individuals with a single breeding queen^20^. Despite gaining widespread interest, the molecular basis of NMR’s unique traits is not thoroughly understood.

Proteome evolution through sequence divergence of proteins has been widely studied^21^. However, the role of other attributes of proteins in the evolution of lineage-specific traits remains underexplored and underappreciated. Proteome remodelling through gain, loss, duplication and fusion of functional protein domains have been widely reported^22–33^. Variation in DNA binding potential has been observed between orthologous transcription factors ^34^. Harris et al and Ramakrishnan et al reported widespread conservation as well as certain lineage specific variations in RNA binding potential of proteins^35,36^. A well-studied structural attribute, the intrinsic disorder, is present in abundance in the hubs of protein-protein interaction networks, epigenetic and transcriptional regulators, signalling proteins, post-translational modification sites, and alternatively spliced exons ^37–45^. The intrinsically disordered regions (IDRs) likely function via couple folding and binding mediated through certain ANCHORs that have increased tendency to fold upon interaction^46–50^ Certain IDRs with suitable pattern of stickers and spacers implicate in phase separation ^51^. A prominent example is that of Heterochromatin Protein-1 (HP1), which forms heterochromatin compartments upon phase separation mediated by an IDR^52^. IDRs are evolutionarily labile^53–55^ and, therefore, can serve as hotbeds for the evolutionary emergence of novel traits.

Despite the wealth of literature on protein intrinsic disorder and the related protein attributes, their implication in the evolution of lineage-specific traits, particularly the extreme ones, has been overlooked. We performed an unbiased study of 18 well curated protein attributes across n orthologous proteins of n mammals. The analyses revealed a high turnover of structural disorder, and concomitantly the order, in naked mole rat specifically. Gain and loss of disorder showed stark functional distinctions. Whereas the former was associated with immune regulation, neurodevelopment, bone growth, apoptosis and tumour suppression, and may function through stabilization of proteins via phase separation, the latter implicated in organogenesis, in particular the heart development and likely function through expression divergence. The disorder turnover was clearly linked to indels, affecting functional sites, domains and redox-sensitivity of proteins, aligning with the stress tolerant physiology of naked mole rat.

## Results

### Evolutionary turnover of intrinsic disorder in Naked Mole Rat

The PCA analysis of 18 protein attributes of orthologous proteins in human, monkey, dog, cat, mouse, guinea pig and naked mole revealed that the intrinsic disorder led the first principal component that explains 17% of total variance (Figure 1A). The second principal component that explains 12 % of variance is led by the protein secondary structures (Figure 1A). We focussed upon intrinsic disorder in the study. An independent PCA analysis of VSL2-predicted intrinsic disorder across species highlighted that the naked mole rat contributes maximally to the leading principal component that explains 23.5 % of total variance (Figure 1B-C). Indeed, the naked mole rat appeared as an outlier in the PCA biplot (Figure 1B). Based on the distribution of variable contributions, we selected top 325 proteins that drove the leading principal component (Figure S1). Out of 325, 171 were negative and 154 were positive loading proteins. Proteins having near zero loadings were enlisted as ‘no change’ genes. Since PCA is not phylogenetically-informed, the observed pattern may result from the phylogenetic relatedness among species. To address this, we performed phylogenetically informed PCA (phylo-PCA), which consistently supported our observations through classical PCA (Figure S2). We further calculated phylogenetically independent contrasts (PICs) for the intrinsic disorder in three categories, by considering phylogeny of naked mole rat, guinea pig and mouse. For the negative loading set, PICs were significantly greater for NMR-G.pig difference when compared with the difference between their common ancestor (x1) and mouse, implying the NMR-specific gain of intrinsic disorder (Figure 1D). Similar trend was observed when compared with the PIC between common ancestor of NMR-G.pig-mouse (x2) and human (Figure 1D). The highest overall magnitude of PICs for NMR-G.pig, followed by x1-mouse and x2-human suggested the NMR-specific divergence of intrinsic disorder (Figure 1D). The trend reversed for the negative loading set, marking loss of intrinsic disorder(Figure 1D). No trend was observed for the no-change set (Figure 1D). Gain of disorder in negative loading set and loss of disorder in positive loading set was further illustrated by plotting distribution of VSL2 disorder scores of proteins across species for each set (Figure 1E). The multivariate analysis, thus, highlighted a significant evolutionary turnover of intrinsic disorder specifically in naked mole rat.

**Figure 1.**
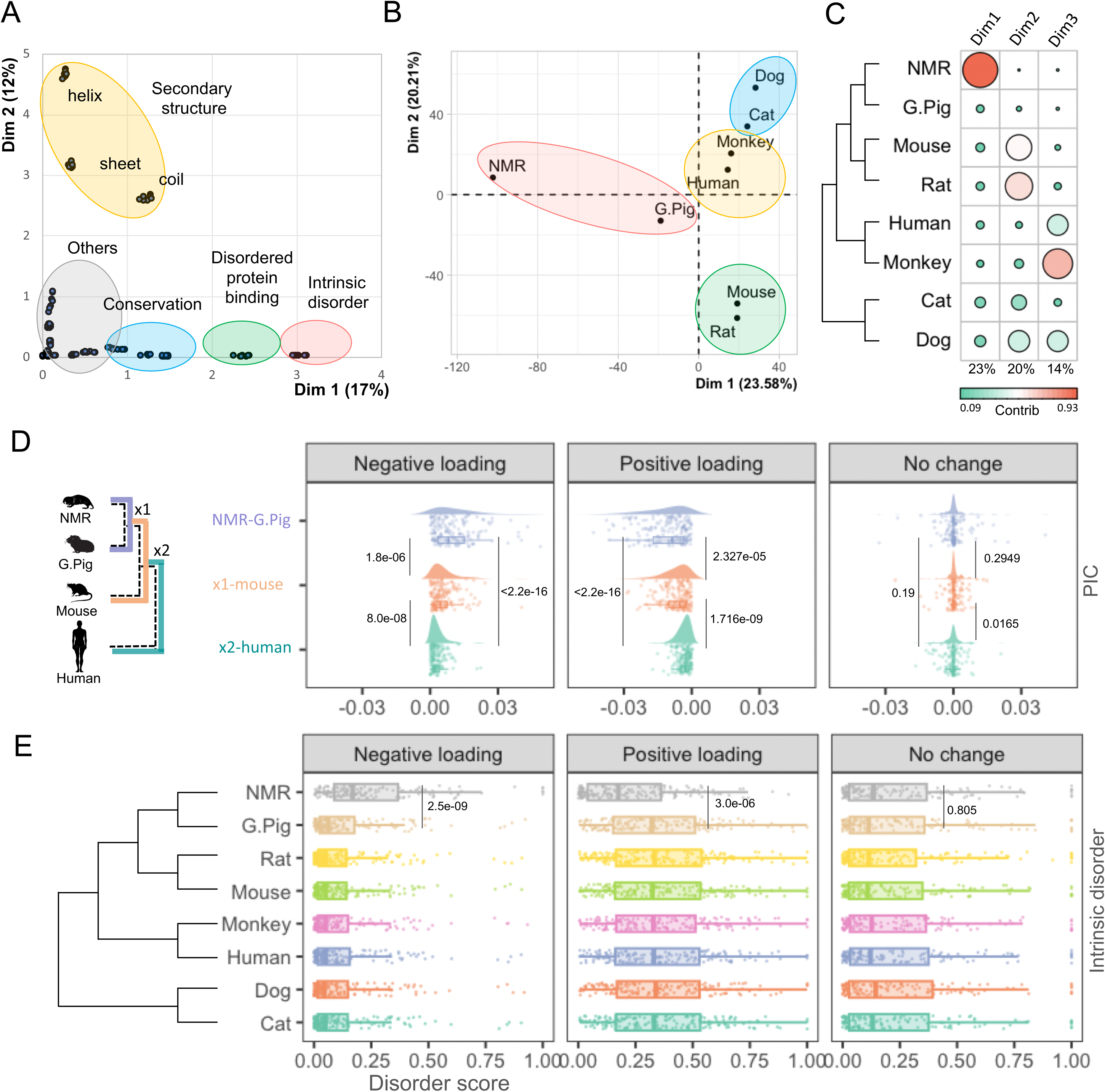
Evolutionary turnover of intrinsic disorder in Naked mole rat. (A) Biplot of contributions of leading principal components of various protein attributes. (B) Biplot of leading principal components of VSL2-predicted intrinsic disorder. (C) Bubble-plot of the relative contributions of each species to the leading components. (D) Distributions of phylogenetically independent contrasts (PIC) values of intrinsic disorder for the following comparisons: NMR – G.pig, x1 (common ancestor of NMR and G.pig) – Mouse, and x2 (common ancestor of NMR, G.pig, and Mouse) – Human. Shown are the cloud-rain plots for negative, no change and positive loading proteins from PC1. (E) Distributions of VSL2-predicted intrinsic disorder for negative, no change and positive lading protein sets. We calculated the p-values using Mann-Whitney *U* tests.

### Functional dichotomy of disorder turnover

Functional enrichment analyses highlighted overrepresentation of proteasomal regulation, immune regulation, neurodevelopment, bone growth, apoptosis, and phosphoinositol related terms in protein set that exhibited gain of disorder (GoD) (Figure 2A). These terms are broadly associated with the distinctive phenotypes of NMR, like longevity, cancer resistance, sustained immune competence, enhanced stress tolerance, skeletal robustness, and neural maintenance throughout lifespan. In contrast, proteins displaying loss of disorder (LoD) had significant enrichment of heart, mesoderm and embryonic development related terms(Figure 2B), tightly aligning with the unique cardiometabolic and cardio-myofilament profile of naked mole rat^56–58^.

**Figure 2.**
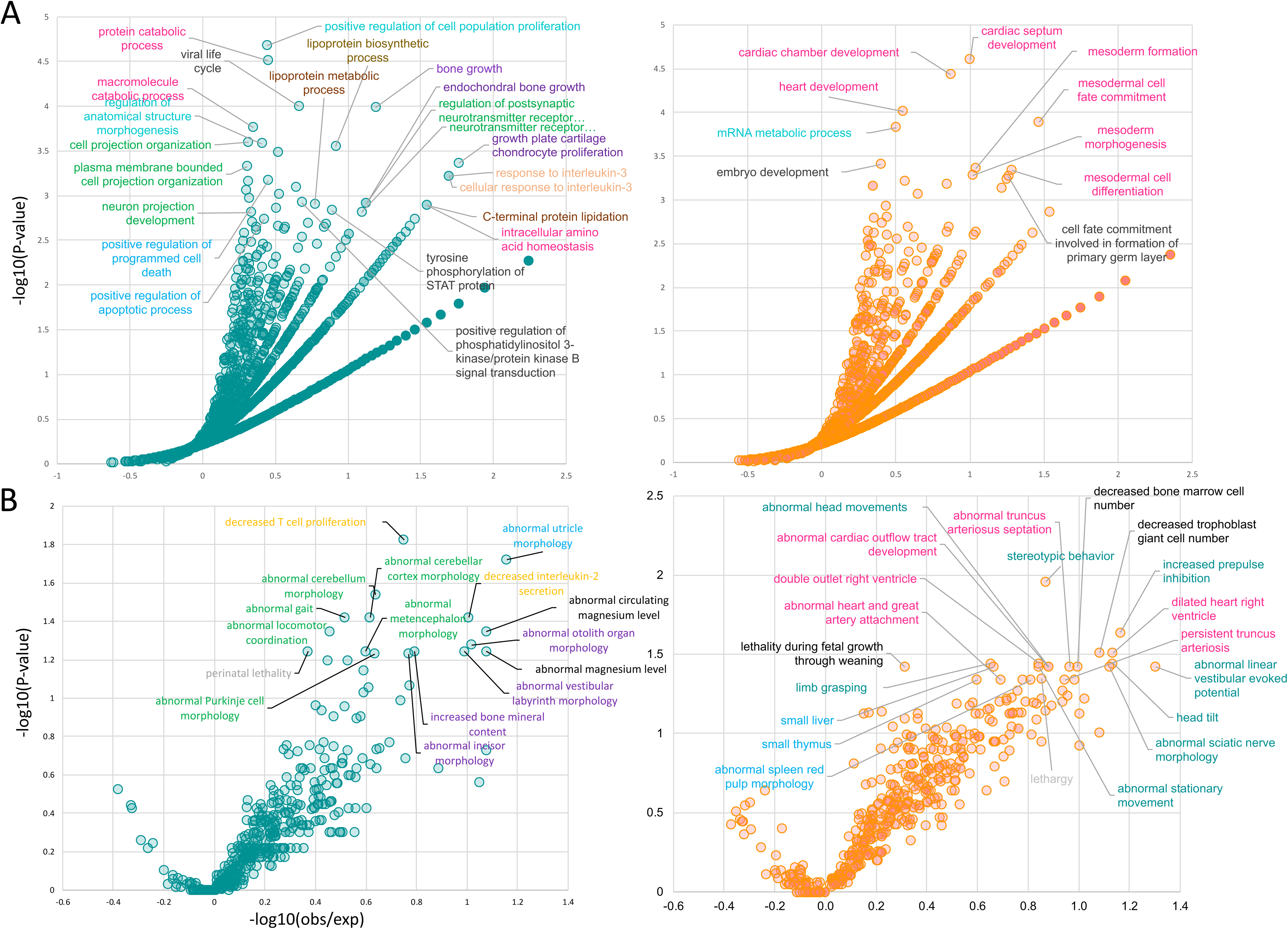
Functional association of proteins exhibiting disorder turnover. (A) Gene ontology enrichment of proteins gaining (left) and losing (right) disorder. (B) Mammalian phenotype ontology enrichment of proteins gaining (left) and losing (right) disorder. Plotted are the FDR values as a function of observed/expected fold enrichment of proteins.

Phenotype enrichment analysis complemented these observations and also revealed additional terms like abnormal erythrocyte magnesium and potassium levels in GoD set, which may relate to unique ionic homeostasis, hypoxia tolerance, and metabolic resilience observed in NMRs (Figure 2C). Conversely, LoD associated phenotypes included trophoblast/placenta development, organogenesis (liver, spleen, thymus), locomotion related terms (Figure 2D), reflecting the NMR’s highly constrained developmental programs, organ maturation, and coordinated motor function necessary for subterranean burrow navigation and eusocial colony living^18,59–65^.

These observations implied functional dichotomy of disorder turnover in NMR. Broadly, the gain of disorder preferentially associate with adaptive, plastic and environmentally responsive traits, while loss of disorder likely safeguards the ordered and constrained developmental process.

### Biophysical consequences of disorder turnover

To assess how the disorder turnover may have globally impacted the biophysical properties of the proteome and consequently the evolutionary adaptations in NMR, we studied the phase separation potential and degradation rates of proteins in each set. Gain of disorder was associated with greater phase separation potential and lower degradation rates (Figure 3A-B). By contrast, proteins losing disorder (LoD) showed lesser phase separation but did not exhibit alteration in degradation rates (Figure 3A-B). These observations imply that the gain of disorder in NMR may have impacted the functional landscape in NMR by influencing the protein stability and functional compartmentalization through phase separation, likely supporting the species’ unique cellular and physiological adaptations.

**Figure 3.**
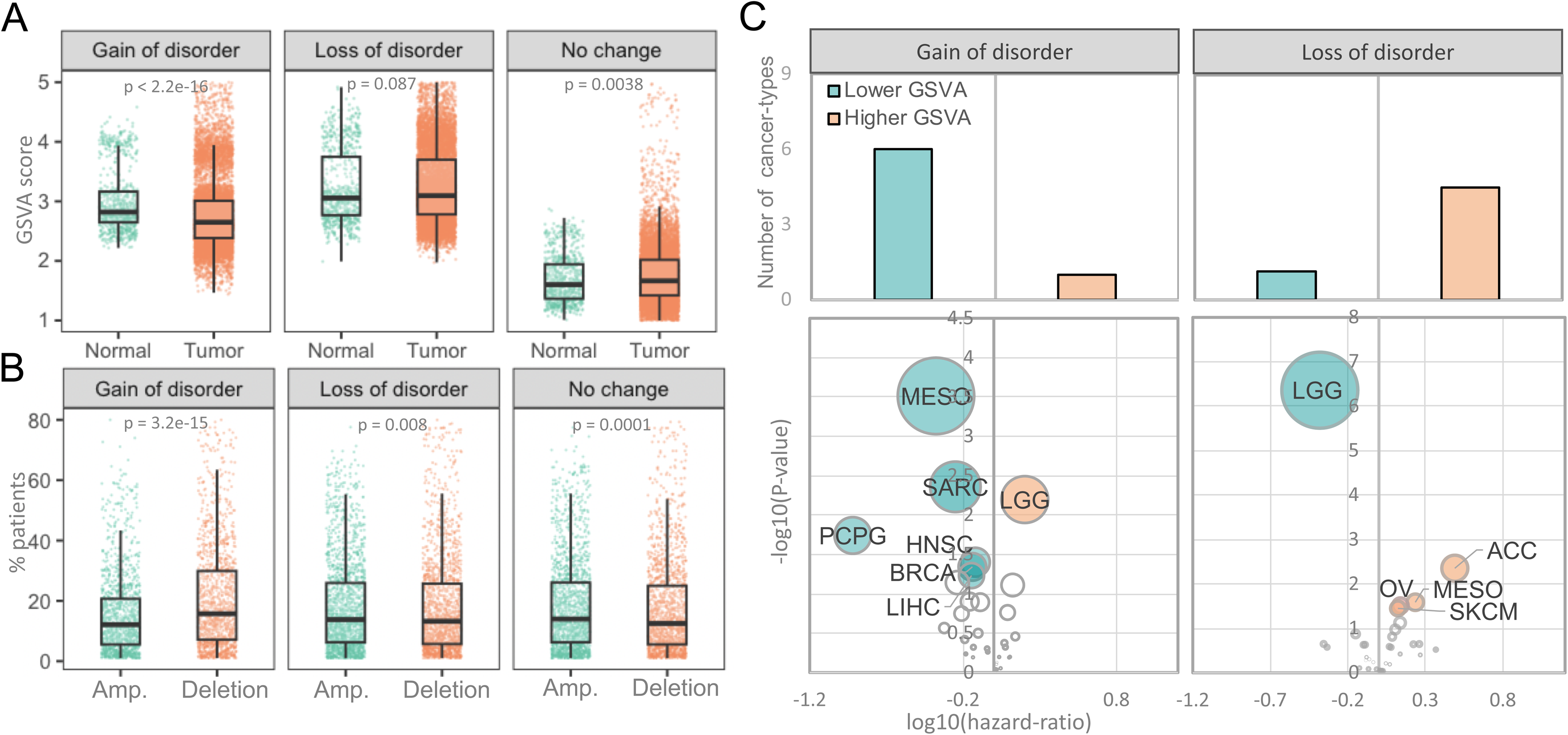
Cancer related properties of proteins exhibiting disorder turnover.(A) Gene set variation analysis (GSVA) scores of genes in ‘gain of disorder’ (negative loading), ‘loss of disorder’(positive loading), and ‘no change’ sets across normal and tumour samples from TCGA. (B) Percentage of TCGA patients exhibiting amplifications and deletions in ‘gain of disorder’ (negative loading), ‘loss of disorder’ (positive loading), and ‘no change’ sets. We calculated p-values using Mann-Whitney *U* tests (C) P-values (negative log2) as a function of Hazard ratio (ratio) for the genes exhibiting low and high GSVA scores in ‘gain of disorder’ and ‘loss of disorder’ sets.

### Gain of disorder associates with anti-tumorigenic properties

Naked mole rat exhibits remarkably lower incidence of cancer^11^. To test the cancer properties of proteins in the GoD and the LoD sets, we performed the gene set variation (GSVA) and copy number variation (CNV) analysis of proteins across cancer and normal samples from TCGA. The GoD set exhibited significantly lower GSVA scores and were often deleted in cancer patients (Figure 4A). LoD set shows no such pattern (Figure 4A). Similarly, there was a greater risk to death, as marked by negative hazard ratios for the lower expression levels of proteins in GoD set (Figure 4B). These observations together implied anti-tumorigenic roles of proteins that GoD set. Coherently, GoD set also exhibited enrichment of functional terms like regulation of apoptosis and phosphoinositol regulation, which are often associated with tumour suppression^66,67^. Among others, we observed the well-known anti-tumorigenic proteins like RAD51C (a DNA repair protein), NQO2 (a stress inducible protector of P53, P63, P33 against degradation), Lumican (a proteoglycan having fas-specific pro-apoptotic role), etc. gained intrinsic disorder in NMR^68–74^. Conversely, the proteins that had lost disorder in NMR did not exhibit such properties (Figure 4). Collectively, these results indicate that gain of disorder in NMR selectively targets proteins involved in cancer suppression, complementing the species’ extraordinary longevity, cancer resistance, and stress tolerance, while loss of disorder preserves core developmental and physiological programs.

**Figure 4.**
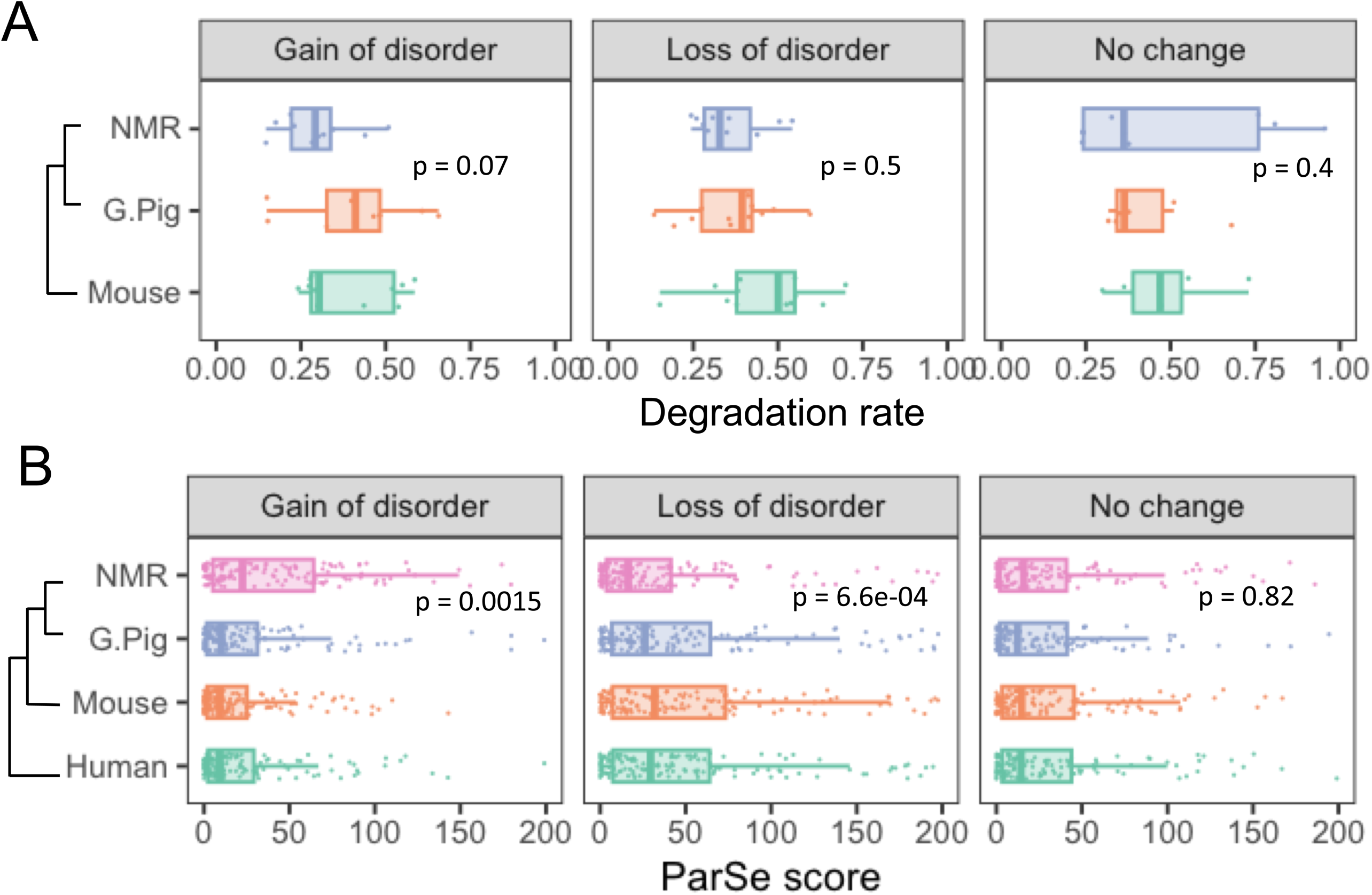
Biophysical correlates of disorder turnover. (A) Protein degradation rates (B) Phase separation potential predicted using ParSe in ‘gain or disorder’, ‘loss of disorder’, and ‘no change’ sets. We calculated the p-values using Mann-Whitney U tests.

### Loss of disorder coincides with the expression divergence

Intrinsically disordered proteins are tightly regulated at transcriptional and translational level. For example, IDPs have lower transcriptional and translational rates and are degraded rapidly when compared with the ordered proteins, implying that the ectopic levels of IDPs may promiscuously interact with other proteins and cause toxicity^75,76^. We, therefore, tested the evolutionary divergence in gene expression of proteins showing disorder turnover in NMR. We calculated expression rates in NMR from guinea pig while taking mouse as an outgroup. Proteins in the LoD set showed significant alteration in expression rates, while the ones in GoD set did not, implying that loss of disorder may allow expression to diverge owing to lower possibilities of promiscuous interactions (Figure 5). Expression divergence of proteins gaining disorder, on the contrary, may lead to abrupt and undesired interactions and, thus, are selected against. The proteins involved in heart development, therefore, lost disorder likely to allow for adaptive expression divergence and tighter regulation in NMR.

**Figure 5.**
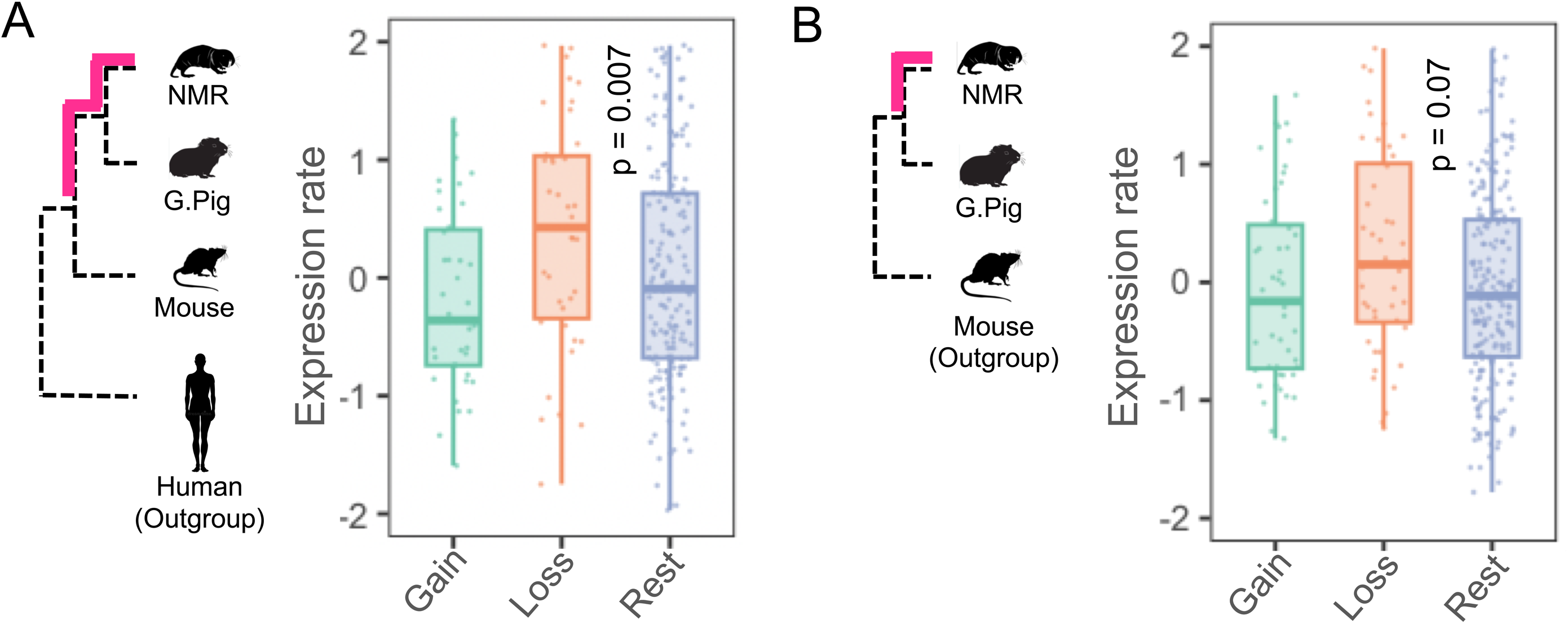
Expression evolution of genes exhibiting disorder turnover. (A-B) Expression rates, calculated through TreeExp framework assuming OU model, (A) by taking NMR as test, mouse as reference, and human as outgroup species, (B) by taking NMR as test, guinea pig as reference, and mouse as outgroup species. We calculated the p-values using Mann-Whitney U tests.

### Disorder turnover is driven by indels affecting functional sites of proteins

To understand the underlying mechanisms driving the disorder turnover in NMR, we performed the multiple sequences alignment of each protein’s ortholog in all the species. By parsing the alignments, we identified the indels (Methods). We first show that indels are the major source of disorder turnover in NMR (Figure 6A-B). Disorder is gained either through insertions of disordered regions or deletion of ordered regions or both (Figure 6A-B). Conversely, disorder is lost through deletions of disordered regions and insertion or duplications of ordered regions (Figure 6A-B). We then tested the representation of functional features like ANCHORs, Pfam domains, short linear motifs (SLiMs), and post-translational modification sites (PTMs) in indels (Methods). The disordered indels (i.e., insertions in GoD set and deletions in LoD set) impacted the gain and loss of ANCHOR predicted binding sites and SLiM motifs, while ordered indels drove the dynamics of Pfam domains (Figure 6C). PTM sites show lesser such distinction (Figure 6C).

**Figure 6.**
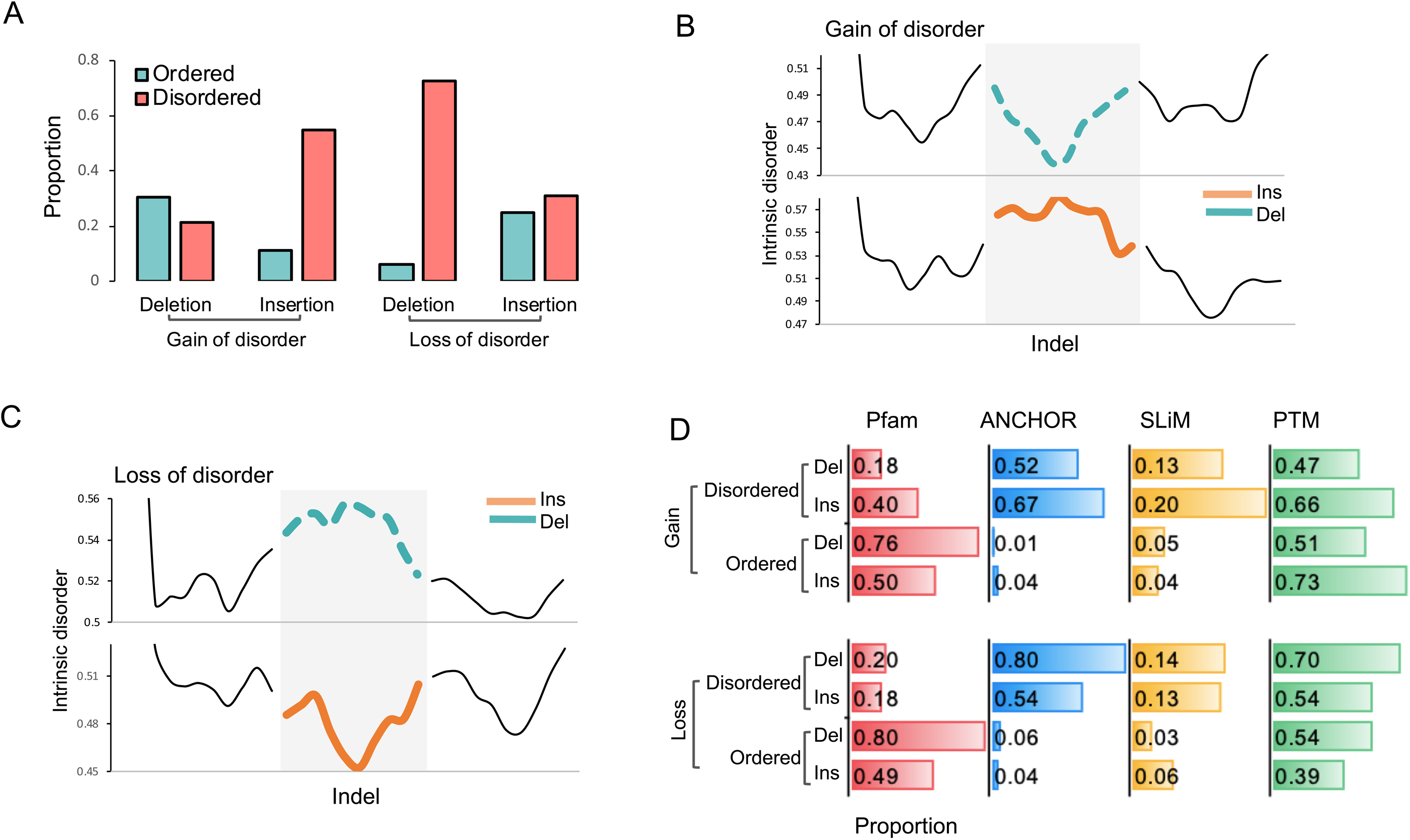
Indel landscape of proteins exhibiting disorder turnover. **(A)** Proportions of insertions and deletions that were ordered or disordered in the ‘gain of disorder’ and ‘loss of disorder’ cases. (B-C). Aggregated disorder scores around indels. For ‘gain of disorder’ in NMR (B), NMR sequences were used, and for ‘loss of disorder’ cases in NMR (C), mouse sequences were used. (D) Proportions of ordered and disordered indels having overlap with Pfam, ANCHOR, SliM and PTMs in gain and loss of disorder cases.

We illustrated several examples of gain and loss of disorder in NMR (Figure 7). The protein FKBP7, FKB10, CAHD1, JAZF1, FBLN5, and TMM3 have gained disorder in NMR, while KLF4, HAND1, 3BP2, HABP4, and TCF20 have lost disorder. FKBP7 and FKBP10 are peptidyl-prolyl cis/trans isomerases (PPIases), known chaperones with multiple functions including protein folding, immune regulation, endoplasmic reticulum stress response, and modulation of signaling pathways^77–79^. FKBP7 had an insertion of a long intrinsically disorder region, while FKB10 had a small deletion that induced structural disorder in the flanking regions in NMR. Gain of disorder in PPIases may modulate conformational flexibility and interaction networks of these chaperones, potentially contributing to enhanced proteostasis and cellular stress resistance in NMR. CAHD1 is an auxiliary regulator of voltage-gated calcium channels that modulates calcium influx and neuronal excitability^80–82^. The gain of intrinsic disorder in CAHD1 may enhance interaction flexibility, potentially contributing to reduced neuronal excitability and resistance to excitotoxic stress in subterranean habitat of naked mole rat. JAZF1 is a key transcriptional repressor in response to metabolic stress^83,84^. It is genetically associated with the diabetes and its deletion causes weight gain and insulin resistance in mice ^85–87^. We show that one of the three zinc fingers is deleted in naked mole rat, which may impact its target selection and binding affinity, likely contributing to the higher insulin sensitivity, low metabolic rates, and consequently the aging in NMR. FBLN5(Fibulin-5) is a glycoprotein that engages in the formation of elastic matrix across several tissues, by interacting with other matrix components like fibrillin-1, tropoelastin, proteoglycan and lysyl oxidases^88–91^. It mitigates the IL-1β induced inflammation of chondrocytes^88^. Fibulin-5 is genetically associated with a wrinkled skin condition ‘Cutis Laxa’ and aging related elastosis in humans and mice ^92–96^. The C-terminal Fibulin domain is important in cellular secretion as well matrix interactions^90^. Interestingly, fibulin domain is partially deleted in FBLN5 of NMR. This may have implication in remodeling of matrix properties in NMR. In particular, it may make the matrix more compliant with the structural properties ascribed by high molecular mass hyaluronic acid, contributing to the elastic wrinkled skin of NMR, and resistance to osteoarthritis^97^. TMM33 is an endoplasmic reticulum (ER) membrane protein that regulates ER morphology and ER–mitochondria contact sites involved in calcium signaling and cellular stress response^98^. Deletion of a centrally located ordered region in NMR’s TMEM33 may weaken or dynamize interactions with ER tethering complexes, potentially limiting excessive ER–mitochondrial calcium transfer and contributing to enhanced cellular stress tolerance

**Figure 7.**
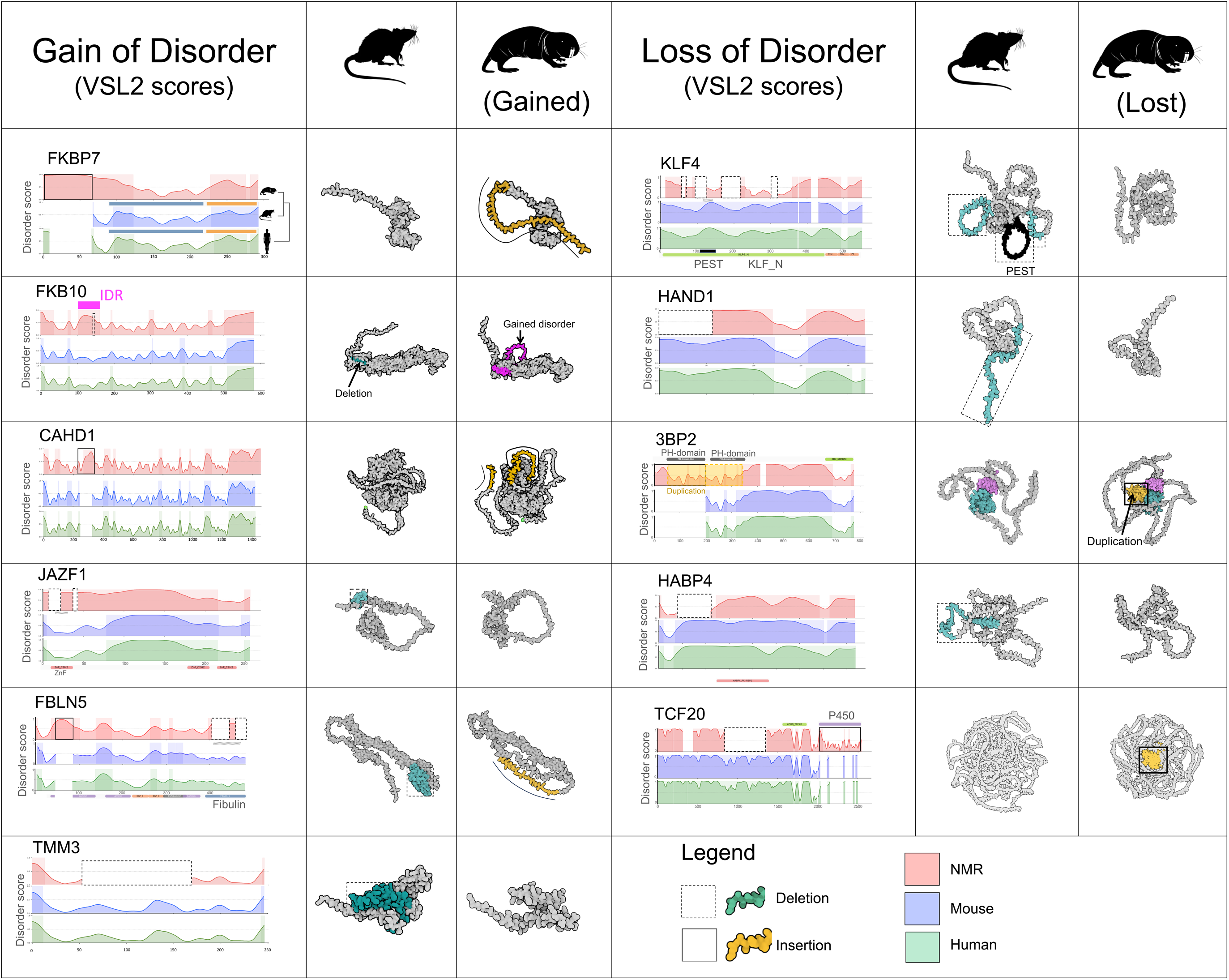
Illustrative examples of evolutionary gain and loss of intrinsic disorder in naked mole rat. Shown are the VSL2-predicted residue-wise disorder scores along the protein sequence alignment of naked mole rat (red), mouse (blue) and human (green) respectively. The insertions and deletions in naked mole rat are highlighted as solid and dashed rectangles. The structural models for mouse and NMR proteins were obtained from AlphaFold3 prediction. Orange colour depicts insertions in NMR and aquamarine colour depicts NMR-deleted regions in mouse.

KLF4 is a transcription factor and epigenetic regulator that acts as a pioneer factor, recruiting chromatin-modifying complexes to remodel chromatin and enabling the reprogramming of differentiated cells to pluripotency as one of the Yamanaka factors^99^. NMR’s epigenome, however, shows remarkable resistance to cellular reprogramming^18^. In NMR’s KLF4, several intrinsically disordered regions, including a PEST motif associated with rapid proteasomal turnover^100^, are deleted, which may increase KLF4 stability and reduce dynamic chromatin remodeling, potentially reinforcing epigenetic stability and contributing to the evolution of reprogramming resistance, longevity, and cancer resistance in the naked mole rat. HAND1 (Heart and neural crest derivatives-expressed protein 1) is essential transcription factor for cardiac morphogenesis, regulating heart tube looping, ventricular specification, and septation during development^101–103^. The naked mole rat exhibits distinctive cardiac physiology characterized by unusually low basal heart rate and contractility but a large cardiac reserve, along with metabolic adaptations that confer exceptional tolerance to hypoxia and ischemic stress^56–58,104,105^. We hypothesize that the loss of a N-terminal intrinsically disordered tail in naked mole rat HAND1 may reduce and interaction promiscuity, potentially stabilizing transcriptional control of cardiac developmental programs. Such structural streamlining may have contributed to the evolution of a cardiometabolic system optimized for energy-efficient function and resilience to chronic low-oxygen environments typical of the species’ subterranean lifestyle. 3BP2 is an adaptor protein having role in phagocytosis and T-cell response. ^106^. NMR had duplication of PH-domain, which interacts phosphatidylinositol 3,4,5-trisphosphate (PIP3) in the plasma membrane ^107^. The double copies of PH domain may increase PIP3 sensitivity and membrane retention, lowering the activation threshold of PI3K signaling and amplifying downstream pathways, aligning with a highly efficient phagocytosis and a unique T-cell response in NMR^108,109^. HABP4 (Hyaluronic Acid Binding Protein 2) is an intrinsically disordered RNA-binding protein involved in mRNA stability, splicing, and post-transcriptional gene regulation, and was originally identified through its ability to bind hyaluronic acid in-vitro^110^. The loss of disorder in HABP4 seems relevant in light of recent evidence showing that naked mole rat cells exhibit elevated RNA synthesis and degradation rates, especially in aging-related pathways, implying a key role of RNA turnover in NMR’s delayed aging^111^. HABP4 also implicates in ribosomal hibernation, a process of keeping the ribosomes in inactive state under stress conditions, thereby contributing to enhanced proteostatsis over decades ^112^. Interestingly, HABP4 is also a known tumour suppressor protein interacting with other tumour suppressor proteins like TP53 and PKC^110,113^. Loss of disorder in HABP4, therefore, is nonrandomly associated with multiple functions that coincide with naked mole rat’s unique physiology.

### Gain of redox-sensitivity in naked mole rat

To gain deeper insight, we calculated the relative amino acid propensities in indels. Deletions were relatively enriched with tyrosine residues, while insertions were marked by tryptophan and cysteine residues (Figure 8A). Redox induced tyrosine modifications, like nitration, contributes to aging and cardiovascular pathologies like atherosclerosis, which naked mole rats are resistant to^58,114,115^. Through inference of tyrosine modifications using a deep learning framework (Methods), we observed their relative enrichment in the deleted regions in NMR (Figure 8B). Their loss may imply greater disease surveillance in in NMR. Tryptophan engages in phase-separation through pi-pi interactions^116^. Their gain may imply increase phase separation and proteostasis in naked mole rat. Cysteine residues can act as redox sensors^117–121^. Overrepresentation of cysteine in inserted regions may hint at gain of redox sensing function in NMR. The observation also aligns with an earlier study, where authors observed abundance of cysteine residues in NMR’s proteome^9^. Through inference of cysteine modifications, we observed relatively greater tendency of cysteine modifications in the inserted regions in NMR (Figure 8B). The role of cysteine is best illustrated by the example of TCF20 protein, which gained an entire P450 domain in naked mole rat (Figure 7). A highly conserved cysteine residue, part of the Phe-X-X-Gly-X-Arg-X-Cys-X-Gly consensus signature in P450 domain, acts as an essential axial ligand to the heme iron^122^. Typically existing as a deprotonated thiolate, the cysteine side chain lowers the iron’s resting redox potential to prevent premature electron transfer^122^. The substrate binding triggers a conformational shift that finetunes the cysteine-iron interaction, sharply increasing the redox potential to “sense” and initiate efficient electron flow for catalysis^122^. These results further warranted the analysis of redox-sensitivity of the observed intrinsic disorder in NMR. Through inference of redox-sensitive disorder^123^, we observed that the redox-sensitive disorder coincided with insertions in naked mole rat when compared to deletions (tested in mouse) (Figure 8C). To control for the overall disorder, the redox-sensitive disorder scores were divided by the overall disorder of the proteins (Methods). We illustrate the example of Calpain-2, a cysteine protease and ANXA6, a calcium-activated membrane-scaffold protein (Figure 8D-E). The injured plasma membrane recruits calcium activated Annexins to form scab-like structures, followed by gradual cleavage of annexins by calpain-2 and finally the release of annexin-containing microvesicles^124^. Calpain-2 showed gain of a cysteine containing disordered region towards N-terminal of cysteine protease domain in NMR, which is critical in sensing the oxidative stress^125^. Concomitantly, ANXA6 has also gained a redox-sensitive an PEST-containing spacer between annexin domains, which may enhance its responsiveness and endow conformationally adaptive binding properties under stress conditions. These observations coherently implied that the gain and loss of intrinsic disorder likely endowed the robustness against redox stress by purging the damaging residues and incorporating the redox sensors.

**Figure 8.**
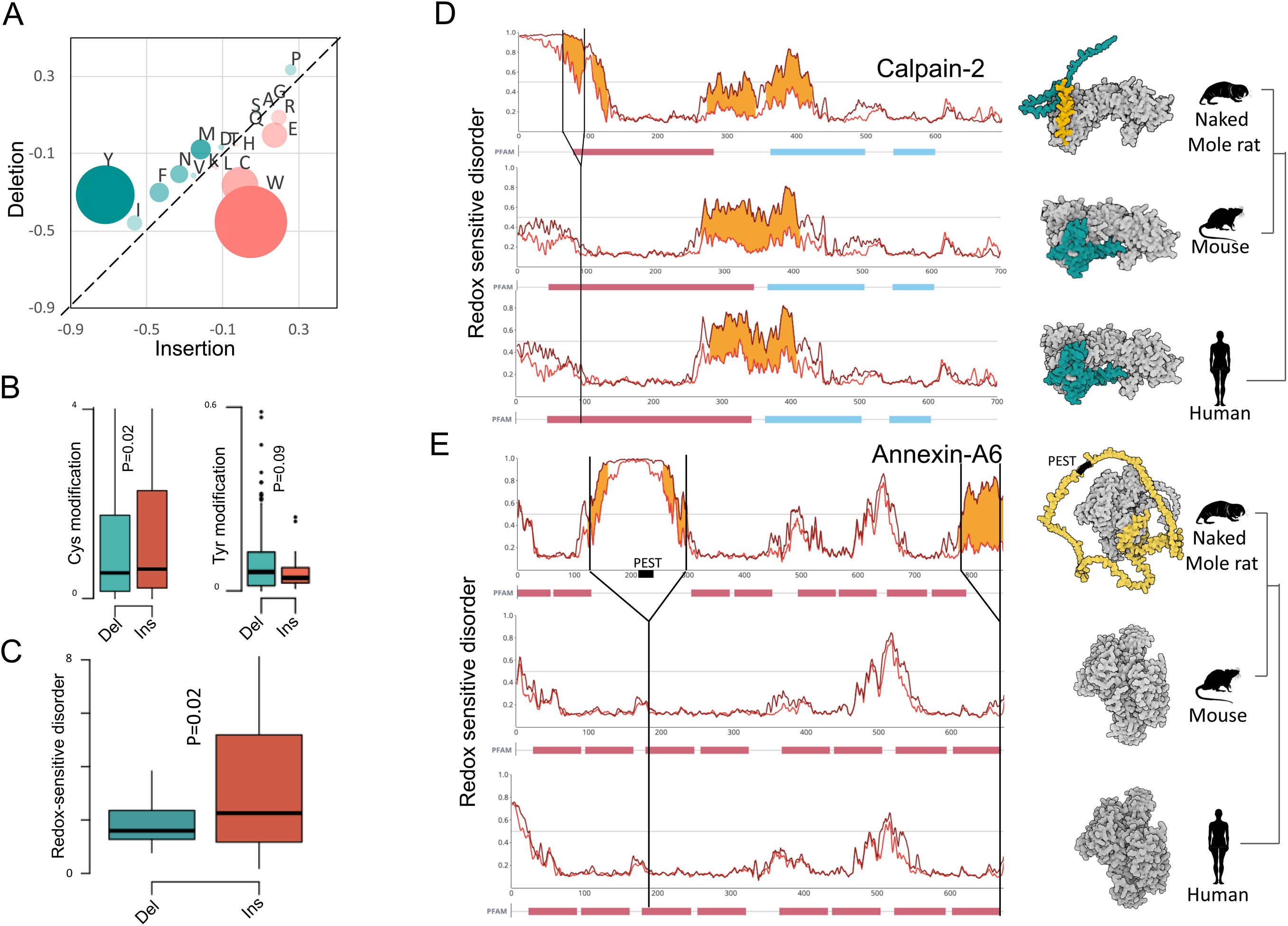
Gain of stress responsive disorder in naked mole rat. **(A)** Scatter plot of amino acid propensities in insertions (tested in NMR) and deletions (tested in mouse), normalized by the respective frequencies of amino acids in the NMR and mouse proteomes respectively. The bubble size are scale based on insertion-to-deletion log fold-change.(B) Inferred Cysteine and Tyrosine modifications in insertions (tested in NMR sequences) and deletions (tested in mouse sequences) in NMR. We calculated p-values using Mann-Whitney U tests. (C) Distribution of redox-sensitive disorder in proteins exhibiting loss and gain of disorder in NMR. The mean of redox-sensitive disorder scores were normalized by overall disorder scores. (D) The illustrative examples (Calpain-2 and Annexin-A6) of gain or redox-sensitive disorder in NMR. Shown are the residue-wise redox-sensitive disorder (brown line), the overall intrinsic disorder (red line), the inferred regions of significant redox sensitivity (orange), and the Pfam domains (bottom) in NMR (top), mouse (middle), and human (bottom). NMR-specific insertions are highlighted through vertical lines. The 3D structures of the proteins modelled using AlphaFold-3 are shown on the right. Orange coloured regions mark the redox-sensitive regions. Aquamarine colour in Calpain-2 marks the N-terminal region. Black colour marks the gained PEST motif in Annexin-A6. (F) Model representing the coordinated roles of Annexins and Calpains in membrane repair, and the role of redox sensitive disorder therein.

## Discussion

Pronounced divergence of intrinsic disorder highlighted an overlooked aspect of naked mole rat biology. The evolutionary gain of disorder correlated with increased phase separation potential and protein degradation rates, aligning with the enhanced proteostasis in NMR. Several studies have demonstrated that intrinsically disordered regions can directly implicate in or facilitate the stress-induced phase separation, a property which is suggested to be adaptive and evolutionarily fine-tuned^126^. It is, therefore, plausible that the gain of disorder may modulate the stress tolerance in NMR by enhancing phase separation potential of protein engaged in NMR. This is supported by the enrichment of stress related functional terms like protein catabolic process, positive regulation of apoptosis, and phophatidylniositol-3-kinase signalling terms among the proteins that gained disorder in NMR. Gained enrichment of cysteine residues, greater overall potential of redox-sensitive disorder in the proteins gaining disorder (tested in NMR) when compared to the ones losing disorder in NMR (tested in mouse), further strengthens the proposal. The examples of Calpain-2 and Annexin-A6 exemplifies the gain of redox-induced disorder in NMR. ANXA6 is a calcium-dependent, pH-sensitive, membrane scaffold that is activated upon hypoxia stress to remodel and repair membrane, coordinate stress signalling and preserve cellular integrity^127,128^. ANXA6 also exhibits tumor-suppressor properties in melanoma, breast cancer, epithelial carcinoma, gastric cancer, and prostate cancer by restraining proliferative and survival signaling (e.g., Ras/MAPK and EGFR pathways) and by promoting membrane and signaling homeostasis^129^. Thus, its upregulation during hypoxia may not only enhance cellular stress tolerance but also limit maladaptive hyperproliferation in chronically low-oxygen environments. Gain of a stress-sensitive intrinsically disordered region between Annexin domains may allow conformationally adaptive membrane properties to sense and interact with varying membrane curvatures under stress conditions. It was noticeable that Calpain-2, which also gained redox-sensitive disorder, also participates in membrane repair by cleaving the annexins after scab formation. Interestingly, gained disordered in Annexin-A6 also had a PEST motif, a well-known substrate of calpain-mediated and other pathways of protein degradation. This implies a complementary and pathway level evolutionary convergence of redox-sensitive intrinsic disorder in NMR (Figure 9).

**Figure 9.**
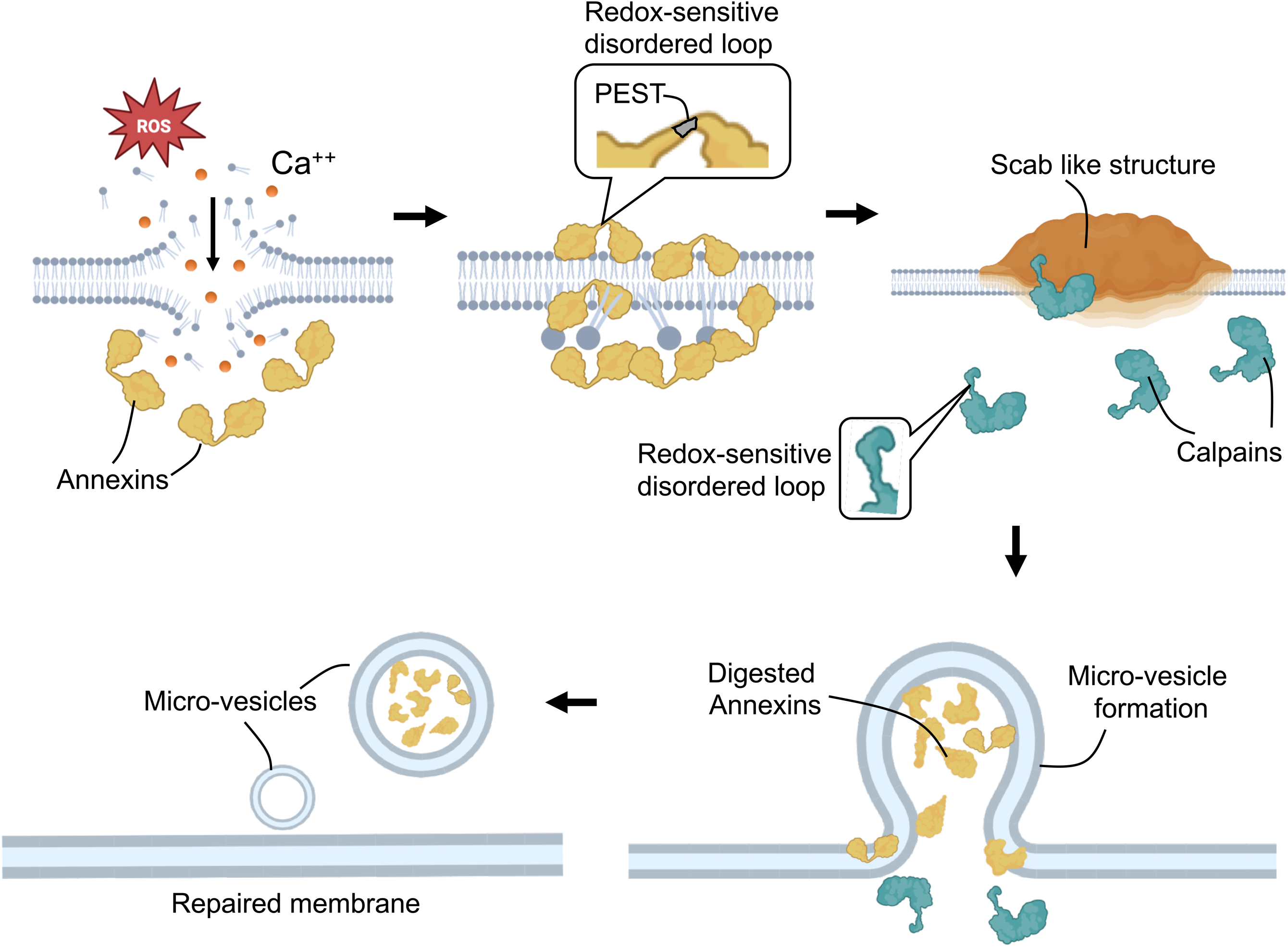
Model illustrating the hypothesized role of intrinsically disordered regions in Annexin A6 and Calpain mediated pathway of membrane repair.

The association between gain of intrinsic disorder and anti-tumorigenic signatures, such as downregulation, recurrent deletions, and negative hazard log-ratios in cancer patients, suggests that enhanced phase-separation propensity and reduced degradation arising from disorder may enable sustained tumour-suppressive signalling in NMR. Through quality-controlled proteostasis, these proteins can buffer cellular stress and support long-term survival under chronically adverse conditions. Loss of disorder in heart development related factors, like HAND1, on the other hand, may imply an evolutionary strategy to endow noise mitigation, avoid promiscuous interactions, and allow for expression divergence. Expression of proteins with intrinsic disorder are often tightly regulated at transcription and as well as translation level in order to avoid inappropriate expression levels, which may cause toxicity. The expression levels of disordered proteins may thus be constrained. Loss of disorder may, however, relieve the constraint and allow the expression to diverge adaptively, likely underpinning its exceptional cardiovascular health, in NMR.

## Conclusion

Altogether, the study highlighted dramatic evolutionary remodelling of naked mole rat’s proteome. The functional polarization of disorder turnover, aligned with the hallmark traits of naked mole rat like stress tolerance, longevity, cancer resistance, immunity, and cardiovascular health underscores its likely adaptive significance. The findings position turnover of intrinsic disorder as dynamic evolutionary tool shaping complex traits. The study also highlights the need to integrate intrinsic disorder into evolutionary framework to understand the molecular basis of lineage-specific innovations.

## Acknowledgement

The authors duly acknowledge the fininacial support from DBT (BT/PR40149/BTIS/137/36/2022, BT/PR40198/BTIS/137/56/2023)

## Methods

### Protein attributes

Protein-wise and residue-wise data on surface accessibility (ASAquick), disordered flexible linker(DFLpred), disorder-mediated protein binding(DisoRDPbind), disorder-mediated RNA binding(DisoRDPbind), disorder-mediated DNA binding (DisoRDPbind), DNA binding (DRNApred), RNA binding (DRNApred), weak conservation (MMseqs2), strong conservation (MMseqs2) MoRFs (MoRFchiBi), helix (PSIPRED), strand (PSIPRED), coil (PSIPRED), protein binding (SCRIBER), signal peptide (SignalP), intrinsic disorder (VSL2B), long disorder (VSL2B) was obtained from a highly curated protein database DescribePROT (https://biomine.cs.vcu.edu/services/webservers-and-databases/). The VSL2B (PONDR) is a multi-feature neural network predictor of intrinsic disorder and is considered to be one of the top performing algorithms in the field^130,131^.

### Orthologues

The protein sequences were obtained from DescribePROT in ‘fasta’ format for all the 8 species. The orthologues of each human protein sequence (reference set) were obtained using OrthoFinder (https://github.com/OrthoFinder/OrthoFinder) *via* an in-house python script. Sequences with one-to-one orthologues were used in the subsequent analyses.

### Principal component analysis (PCA)

We used ‘PCA’ function of ‘FactoMineR’ r-package (https://github.com/cran/FactoMineR/) to perform PCA. Data was Z-scaled prior to PCA. First PCA was performed on the complete data covering all protein attributes across all species. Second PCA was performed on VSL2-predicted intrinsic disorder across all species. The ‘var$coord’ values in the second PCA were taken as ‘loadings’. Loading of variable j on prinpal component k is given by,

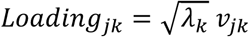

Where *λ_k_* is the eigen value of *k^th^* principal component, and *v_jk_* is the *j^th^* element of the eigen vector corresponding to *k^th^* principal component.

Contribution of *j^th^*variable to *k^th^* component was obtained using:

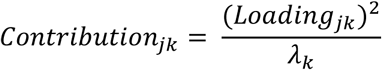

Based on the threshold determined by plotting the distribution of ‘contribution’ values, 171 proteins were categorized as negative, and 154 proteins as positive loading proteins.

Phylogenetically informed PCA was performed using ‘phyloPCA’ package (https://github.com/mwpennell/phyloPCA)

### Phylogenetic independent contrasts (PICs)

Phylogenetic independent contrasts calculates normalized difference of trait values between the sister lineages in the phylogenetic tree under the assumption of Brownian evolution. PICs are statistically independent and allow for appropriate evolutionary interpretation, which otherwise is affected by shared ancestry of lineages. We used ‘pic’ function in ‘phytools’ r-package (http://www.phytools.org/) to correctly infer the evolutionary gain and loss of intrinsic disorder in NMR.

### Protein degradation rates and phase separation potential

Protein degradation rates in NMR, guinea pig, and mouse were obtained from Swovick et al^132^. Phase separation potential was calculated using ParSe algorithm (https://stevewhitten.github.io/Parse_v2_web/).

### Cancer related properties

TCGA cancer and normal data was mined using GSCA pipeline (https://guolab.wchscu.cn/GSCA/<u>#/</u>). For gene set variation analysis (GSVA) and CNV (heterozygous) analysis, results for all cancer types were collated together. GSVA transforms gene expression data into sample-wise enrichment scores, enabling direct comparison of our gene set between cancer and normal samples. Unlike gene centric methods, GSVA is less sensitive to noise and robustly captures the variation of pathway activity over a sample population in an unsupervised manner, We used categorical hazard ratio in log10 scale for the higher (above median) and lower (below median) GSVA scores. We only considered ‘overall survival’ for this analysis.

### Evolutionary rates of gene expression

The expression rates were calculated using ‘TreeExp’ R-package assuming OU model of transcriptome evolution (https://github.com/hr1912/TreeExp). Given a set of orthologous genes in a test (t), a reference(r) and an outgroup species(o), TreeExp calculates expression distances *d_tr_, d_to_, d_ro_* under OU model and considers following statistics, *d_tr_=d_to_-d_ro_*assuming the additivity of expression distances. The *d_tr_* scores are then converted to z-scores for statistical inference.

TPM values of RNA-seq data of brain, heart, kidney, liver, and skin for human, mouse, guinea pig and naked mole rat were obtained from ‘bgee’ (https://www.bgee.org/) and log2-transformed. We made bins of 50 proteins from the list sorted on loadings (positive to negative) . For each bin expression rate was calculated using NMR as test, mouse and reference, and human as an outgroup species. Top 10 (positive loading, loss of disorder), bottom 10 (negative loading, gain of disorder) bins were compared with rest of the bins. In another comparison, NMR was taken as test, guinea pig as reference, and mouse as an outgroup species.

### Identification of Indels

Multiple sequence alignments of orthologous proteins of 8 species were performed using MAFT (https://mafft.cbrc.jp/alignment/software/linux.html) with default parameters. We wrote python scripts to fetch NMR specific indels with minimum length of 3 residues. An indel at locus *i* is defined as NMR-specific if,

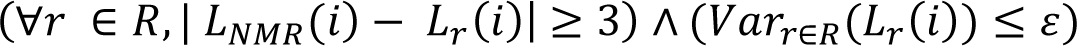

where *i* denotes an alignment block, *L_NMR_(i)* is the aligned sequence length at locus *i* in naked mole rat, *L_r_(i)* is the aligned sequence length at locus *i* in reference species *r*, *R={guinea pig,mouse,human,chimpanzee,cat,dog}* is the set of rest of the reference species, *∀_r_∈R* indicates that the condition holds for all reference species, |*L_NMR_(i)−L_r_(i)*| *≥* 3 enforces a minimum length difference of 3 aa between NMR and each reference species (capturing an insertion or deletion), *Var_r∈R_(L_r_(i))* is the variance of sequence lengths across the reference species at locus *i*, and *ɛ* is a small positive tolerance parameter allowing minor variation among reference species. The residue-wise VSL2 disorder scores were mapped onto multiple sequence alignments using inhouse python script.

### Mapping functional sites

We used standalone ‘hmmscan’ function of HMMER package (http://hmmer.org/download.html) to scan the sequences against Pfam-A database. API of AIUPred v2(https://aiupred.elte.hu/) was used to predict the ANCHOR binding regions. Short-linear motifs (SLiMs) were extracted running the script of SLIMFindeR (https://github.com/vitkl/SLIMFinderR). PTM sites obtained using NetPhos-3.1 (https://services.healthtech.dtu.dk/services/NetPhos-3.1/) .

### Redox sensitivity

Amino acid propensities of indels were calculated by counting each amino acid and dividing by the length of indel. AA propensities were normalized by overall frequencies of amino acids in NMR (for insertions) and mouse (for deletions) proteomes. Log10 transformed values were plotted.

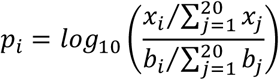

We inferred Cysteine S-nitrosylation, S-palmitoylation, S-sulfhydration, S-sulfenylation, S-sulfinylation using deep learning framework of pCysMod (http://pcysmod.omicsbio.info/). Tyrosine nitration was interred using deep neural networks of deepNitro (https://deepnitro.renlab.org/). Distribution of mean scores of indels were plotted.

We calculated the Redox-sensitrve disorder using the API of AIUpred v2 (https://aiupred.elte.hu/) with the parameter ‘analysis_type’ as ‘redox’. AIUpred infers redox sensitivity of disorder by calculating the interaction energies under reducing and oxidizing conditions. The method considers altering the cysteine’s contribution to interaction energies in order to capture the stabilizing effect of disulfide bonds under oxidizing conditions. Residues exhibiting substantial difference in the interaction energies between reduced and oxidized states are classified as redox-sensitive. The mean redox-sensitive disorder scores were normalized by the overall disorder scores of proteins for statistical comparisons.

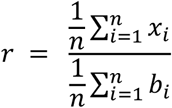

Where *x_i_* is the redox sensitive disorder score and *b_i_* is the overall intrinsic disorder scores of *ith* residue of the protein. We plotted the residue-wise maps of redox-sensitive disorder using AIUpred web interface.

### 3D structure modelling

AlphaFold-3 with default parameters was used to model 3D structures and associated Predicted Aligned Error (PAE) matrices and of the proteins in the respective species. ‘Spacefill’ illustrative cartoons with the region of interest coloured using selection helpers were drawn on AlphaFold-3 interface.

### Statistical tests of significance

We did not assume any underlying distribution of our datasets and, therefore, implemented non-parametric test of significance, namely Mann-Whitney *U* test using ‘wilcox.test’ function in R. The enrichments of gene ontology and mammalian phenotype ontology terms were tested using hypergeometric tests followed p-value corrections for multiple testing using Benjamini Hochberg method on Toppgene (https://toppgene.cchmc.org/enrichment.jsp) and ModPhEA (https://evol.nhri.org.tw/phenome2/) servers respectively.

---

**Figure S1.**
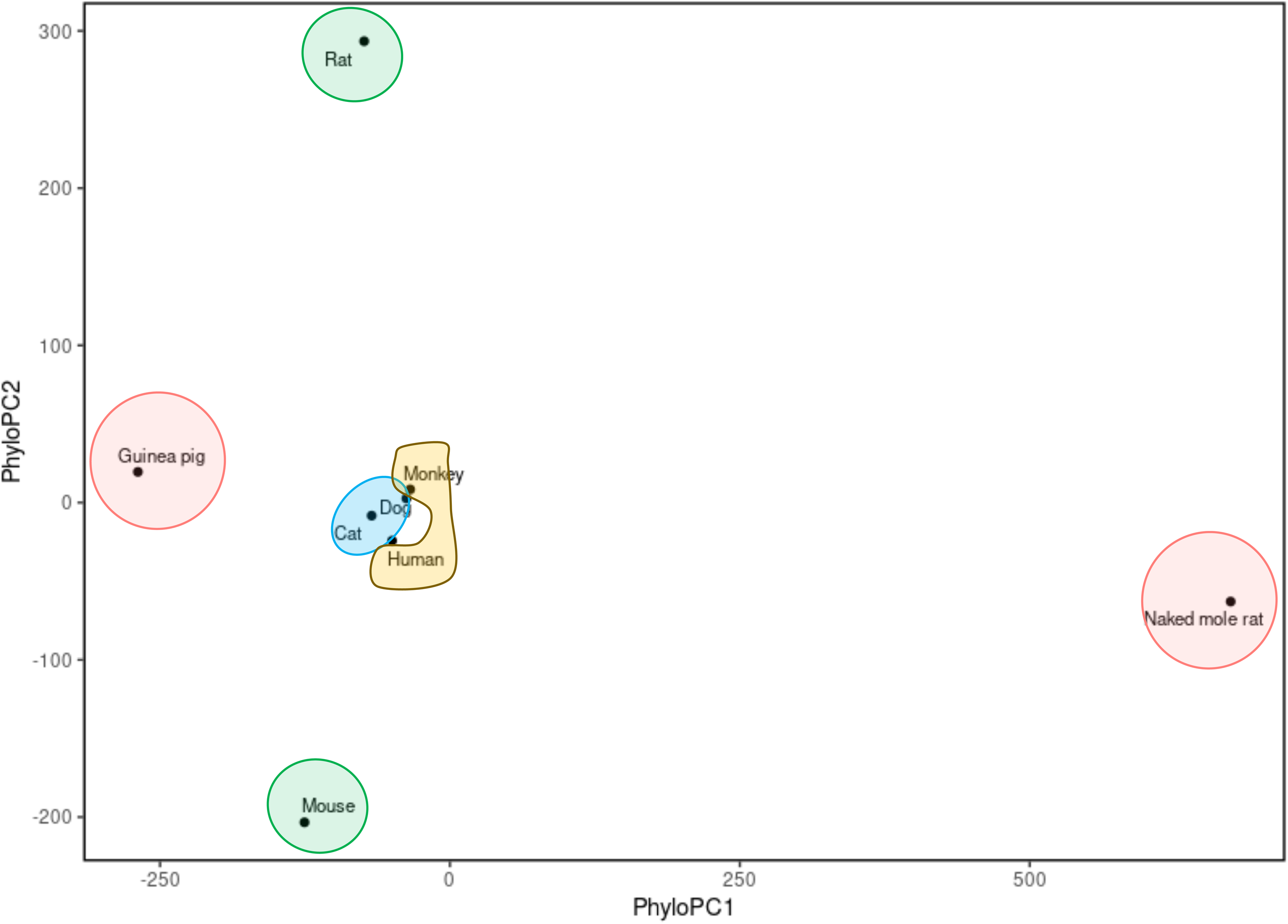

**Figure S2.**
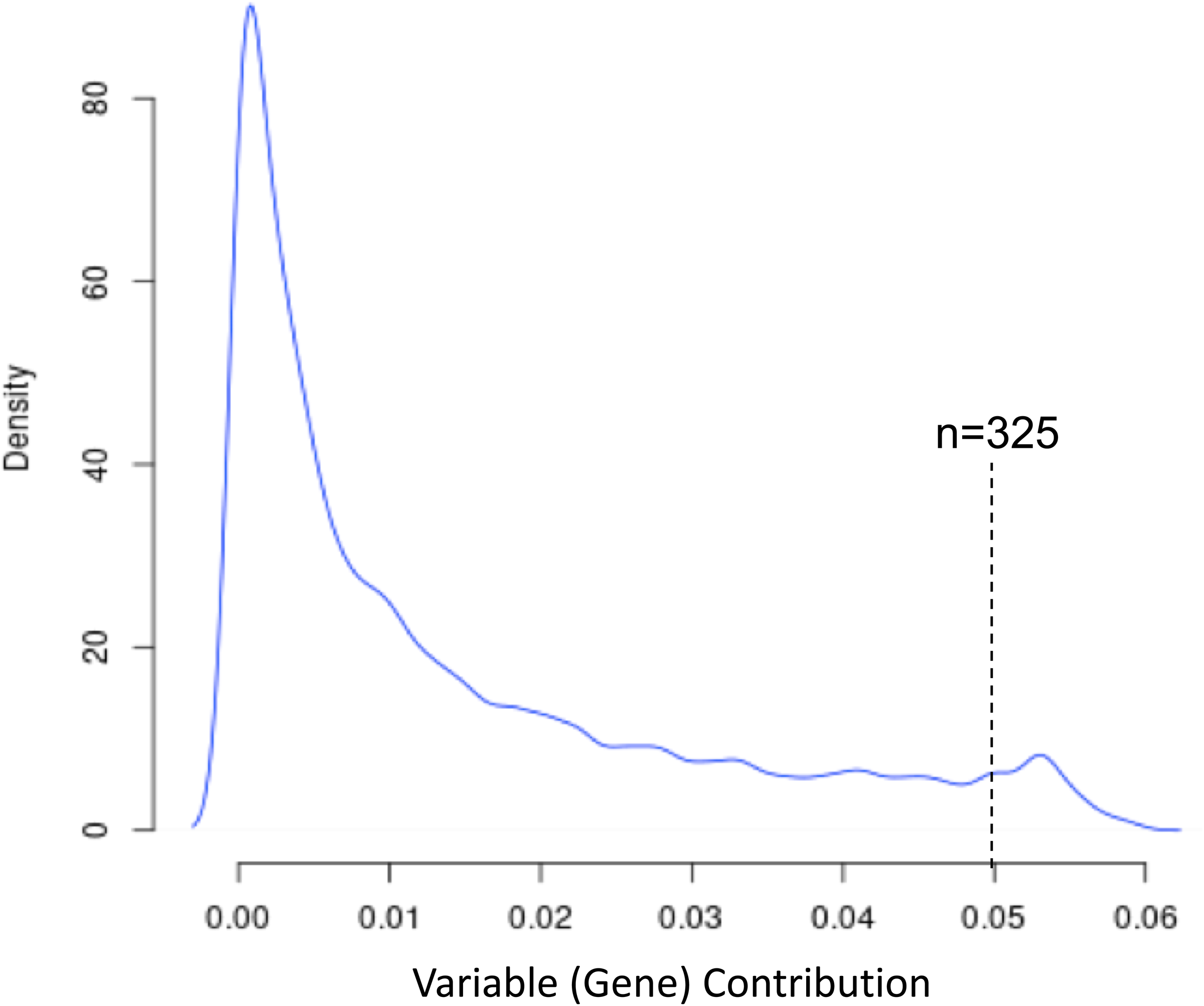

## Notes

### Competing Interest Statement

The authors have declared no competing interest.

